# Idiosyncrasy and generalizability of contraceptive- and hormone-related functional connectomes across the menstrual cycle

**DOI:** 10.1101/2024.10.24.620112

**Authors:** Katherine L. Bottenhorn, Taylor Salo, Emily G. Jacobs, Laura Pritschet, Caitlin Taylor, Megan M. Herting, Angela R. Laird

**Affiliations:** Department of Population and Public Health Sciences, University of Southern California, Los Angeles, CA, USA; Department of Psychology, Florida International University, Miami, FL, USA; Lifespan Informatics and Neuroimaging Center (PennLINC), Department of Psychiatry, Perelman School of Medicine, University of Pennsylvania, Philadelphia, PA, USA; Neuroscience Research Institute, University of California, Santa Barbara, CA, USA; Department of Psychological and Brain Sciences, University of California, Santa Barbara, CA, USA; Department of Physics, Florida International University, Miami, FL, USA; Center for Imaging Science, Florida International University, Miami, FL, USA

**Author notes:** Corresponding Author*: Dr. Katherine L. Bottenhorn, Postdoctoral scholar, Keck School of Medicine of USC, University of Southern California, Los Angeles, CA, USA |.

**Keywords:** functional connectivity, estradiol, progesterone, hormonal contraceptives, menstrual cycle

## Abstract

Neuroendocrinology has received little attention in human neuroscience research, resulting in a dearth of knowledge surrounding potent and dynamic modulators of cognition and behavior, as well as brain structure and function. This work addresses one such phenomenon by studying functional connectomics related to ovarian hormone fluctuations throughout the adult menstrual cycle. To do so, we used functional magnetic resonance imaging (fMRI) and hormone assessments from two dense, longitudinal datasets to assess variations in functional connectivity with respect to endogenous and exogenous endocrine factors throughout the menstrual cycle. First, we replicated prior findings that common, group-level and individual specific factors have similar relative contributions to functional brain network organization. Second, we found widespread connectivity related to hormonal contraceptive (HC) use, in addition to sparser estradiol- and progesterone-related connectivity, and differential generalizability of these subnetworks that suggests progestin-specific impacts on functional connectivity in HC users. These results provide novel insight into within-individual changes in brain organization across the menstrual cycle and the extent to which these changes are shared between individuals, illuminating understudied phenomena in reproductive health and important information for all neuroimaging studies that include participants who menstruate.

**Author Summary:** Endocrine modulation of brain function across the menstrual cycle is poorly understood. Human neuroimaging research on the menstrual cycle has long relied on group- and or coarse, within-individual cycle stage-differences, overlooking considerable individual differences in brain organization, the menstrual cycle, and hormone concentrations. Here, we take a multi-dataset approach to identify idiosyncratic contraceptive- and hormone-related functional connectivity from within-individual neuroendocrine dynamics and then test the generalizability of this connectivity to other individuals. In doing so, we identified idiosyncratic hormone-responsive subnetworks that are somewhat generalizable to other individuals, though this generalizability is complicated by hormonal contraceptive use, potentially reflecting differential connectivity between contraceptive formulations. Thus, this work illuminates individual similarities and differences in neuroendocrine dynamics across the menstrual cycle.

## Introduction

As a critical node of the endocrine system, the brain influences and is influenced by peripheral hormone fluctuations, which can impact its structure and function (Albert et al., 2017; Beltz & Moser, 2020; Brown et al., 2015; Galea et al., 2017; Taylor et al., 2020). A prime example of this bidirectional relationship comes from fluctuating ovarian hormones, (e.g., estradiol and progesterone) across the menstrual cycle, which are integral to the female reproductive system. Estradiol influences dendritic spine density (Hao et al., 2006), major neurotransmitter systems (Barth et al., 2015; Bendis et al., 2024; Diekhof, 2018; Zsido et al., 2017), learning and memory (Jacobs et al., 2016; Jacobs & D’Esposito, 2011; Taxier et al., 2020), and emotion regulation (Graham et al., 2017; van Wingen et al., 2011). Conversely, progesterone’s role in cognition is largely inconclusive (Henderson, 2018), though it is related to significant fluctuations in medial temporal lobe volume (Taylor et al., 2020; Zsido et al., 2023). Hormonal contraceptives (HCs) with synthetic estrogens and progestins disrupt this cycle. Further, HC use has been linked to increased risk of depression (Skovlund et al., 2016); altered gray matter volume in medial temporal, PFC, and hypothalamic regions (Brønnick et al., 2020; Chen et al., 2021); and differences in cognitive function (Brønnick et al., 2020; Warren et al., 2014).

However, endocrine factors have received little attention in human neuroscience in either sex (Taylor et al., 2021; Will et al., 2017). Of the more than 50,000 published human MRI studies, only 0.5% address sex hormones or other endocrine factors (Jacobs, 2023b; Taylor et al., 2021). Here, we focus on ovarian hormone fluctuations during the adult menstrual cycle: a period of 25–30 days, during which estradiol levels undergo ∼8-fold changes; progesterone levels, ∼80-fold changes. Open questions remain regarding the roles of these daily and weekly hormone fluctuations in brain function, in addition to the influences of HC use (Pletzer & Kerschbaum, 2014). Much of the extant literature focuses on contrasts between menstrual cycle phases, which obfuscate hormone dynamics, as endogenous estradiol and progesterone fluctuations are not characterized by discrete transitions between menstrual cycle phases, but curvilinear progressions across the cycle. Within the remarkably undersized proportion of neuroimaging papers that address endocrine factors, fewer than 8% of published human neuroimaging studies of the menstrual cycle have assessed individuals at more than three time points and only 30% assessed individuals more than twice (Dubol et al., 2021). Furthermore, menstrual cycles vary within an individual as much as they do between individuals (Fehring et al., 2006; Jasienska & Jasienski, 2008; Stern et al., 2022). Studies of exogenous hormone manipulations, on the other hand, suggest HC use creates a hyperprogestogenic state in the brain characterized by altered medial temporal, prefrontal, and parieto-occipital connectivity (reviewed by (Casto et al., 2022)). However, extant studies of both endogenous and exogenous hormones across the menstrual cycle focus largely on between-group differences and are unable to model or estimate individual differences, or idiosyncrasies, in these effects. Thus, the attention paid to ovarian hormones and endocrine rhythms in the extant human neuroimaging literature is insufficient to characterize neuroendocrine dynamics across the menstrual cycle. Given the expression of ovarian hormone receptors across the brain *and* the interactions of ovarian hormone metabolites with major neurotransmitter systems (Barth et al., 2015; Baulieu et al., 1996; Brinton et al., 2008), these fluctuations have large, currently overlooked consequences for brain function.

Deep phenotyping and precision neuroscience studies can address this gap by densely sampling individuals with repeated measurements over days, weeks, and months to better understand both between- and within-individual variability in the brain (Gordon et al., 2017; Jacobs, 2023a; Poldrack, 2017; Poldrack et al., 2015; Pritschet et al., 2021; Pritschet & Jacobs, 2024). Deep and/or dense phenotyping approaches allow for more precise measurement and estimation of previously unmodeled sources of individual differences. Gratton et al. (2018) found that common group factors and stable individual-specific factors have near-equal contributions to similarity of functional brain networks, illuminating the balance of generalizability and idiosyncrasy in functional connectomics. While the same work found that within-individual differences over time are responsible for a relatively small proportion of network similarity, deep phenotyping of the human brain across the menstrual cycle shows notable changes in medial temporal lobe structure, in addition to cortical and cerebellar functional network organization (Fitzgerald et al., 2020; Pritschet et al., 2020; Taylor et al., 2020). While dense-sampling approaches offer a nearly unparalleled opportunity to better elucidate hormone–brain interactions, only a minority of such studies include endocrine assessments (Arélin et al., 2015; Barth et al., 2016; Filevich et al., 2017; Pritschet et al., 2020; Zsido et al., 2023). Here, we use three such datasets to investigate roles of ovarian hormones in brain network organization across the menstrual cycle. The first two are from the 28andMe projects, which include daily fMRI, hormone, and behavioral assessments over two 30-day periods in a single premenopausal woman while naturally-cycling (28andMe; (Pritschet et al., 2020)) and again using hormonal contraceptives (28andOC) after a 10-month wash-in period. The third is from the Dense Investigation of Variability in Affect (DIVA) Study (Bottenhorn and Salo et al., 2024), which includes weekly fMRI, hormone, and behavioral assessments over 2–5 weeks in three premenopausal women: one naturally cycling and two using hormonal contraceptives. These data present an opportunity to study variability in the brain related to the menstrual cycle and to better understand neuroendocrine dynamics affecting over 50% of the population.

The present study examines the roles of HC use and ovarian hormone concentrations in functional brain network organization across the menstrual cycle. First, we aimed to replicate the findings of Gratton et al. (2018) regarding the degrees of generalizability and idiosyncrasy in functional network organization within our multi-dataset sample and extend this approach to assess network similarity between different datasets and HC use. To do this, we calculated the similarity (i.e., product-moment correlation) between functional brain networks across each category (i.e., group, dataset, HC use, individual). Second, we identified roles of HC use and ovarian hormones in the brain’s functional connectome by a combination of graph theory and machine learning. To this end, we used the network-based statistic in a machine-learning framework (NBS-Predict; (Serin et al., 2021; Zalesky et al., 2010)) to identify HC- and hormone-sensitive subnetworks in the 28andMe and 28andOC datasets. We then assessed how these subnetworks generalized to the independent DIVA dataset. This approach leverages a multivariate, predictive connectomics approach that is more generalizable and robust than univariate, descriptive approaches (Mantwill et al., 2022; Marek et al., 2022; Shen et al., 2017). We expected that dataset- and individual-specific features would dominate functional connectivity, compared with HC use, and that brain regions demonstrating high concentrations of estrogen receptors (e.g., prefrontal cortex, hippocampus and broader medial temporal lobe; (Wang et al., 2010)) would be involved in hormone-sensitive subnetworks over the course of the menstrual cycle. Investigating the interplay of endocrine factors in functional connectomics on daily and weekly scales will improve our understanding of the brain, contribute to a growing literature regarding neuroendocrine dynamics in both sexes (Ballard et al., 2022; Barth et al., 2016; Coenjaerts et al., 2023; De Filippi et al., 2021; Dubol et al., 2021; Fitzgerald et al., 2020; Greenwell et al., 2023; Grotzinger et al., 2024; Hidalgo-Lopez et al., 2021, 2023; Liparoti et al., 2021; Martínez-García et al., 2021; Mueller et al., 2021; Pritschet et al., 2020; Taylor et al., 2020; Zsido et al., 2023) and advance knowledge of differential health outcomes for women and people who menstruate.

## Methods

Here, we leverage three complementary datasets to assess how hormones and behavior are associated with brain network variability: 28andMe, 28andOC, and DIVA.

### Participants and Study Design

#### 28andMe, 28andOC

Data were collected from one premenopausal, female participant (monikered “sub-01”; age 23, right-handed) with no history of neurological or psychiatric diagnoses, a history of regular menstrual cycles, and no hormonal contraceptive use in the prior year. She completed daily hormone, behavioral, and MRI assessments for 30 days (28andMe) and then again for 30 days one year later (age 24), while using hormonal contraceptives (0.02 mg ethinyl-estradiol, 0.1 mg levonorgestrel, Aubra, Afaxys Pharmaceuticals) started 10 months prior (28andOC). Written, informed consent was obtained from this participant before data collection began, in accordance with the University of California, Santa Barbara Human Subjects Committee.

#### DIVA

Data were collected from three premenopausal, female participants (monikered “sub-02”, “sub-03”, and “sub-04”; age range = 26-31 years). At the time of data collection, two participants were using hormonal contraceptives for at least 6 months and reported no endocrine conditions (sub-04: 0.035 mg ethinyl-estradiol, 0.025 mg norgestimate, Feymor, Amneal Pharmaceuticals; sub-03: 0.02 mg ethinyl-estradiol, 1 mg norethindrone acetate, Blisovi Fe) and the third (sub-02) was freely cycling, with a history of regular menstrual cycles, and had not used hormonal contraceptives in the prior year. Participants completed behavioral assessments and collected saliva samples twice a week, 3–4 days apart, and completed MRI scanning sessions once a week (on a behavioral + hormone collection day). Written, informed consent was obtained from each participant before data collection began, in accordance with Florida International University’s Institutional Review Board approval.

### Endocrine Data

#### 28andMe, 28andOC

Endocrine measures include serum assessments of gonadotropins (luteinizing hormone; LH and follicle stimulating hormone; FSH) and sex steroid hormones 17*β-*estradiol and progesterone concentrations ([E_2_] and [P_4_], respectively). Blood samples (10-mL) were collected daily at or within 30 minutes of 10:00 AM local time. After clotting at room temperature for 45 min and centrifugation at 2,000 g for 10 min, samples were then aliquoted into three 1-mL sterile cryovials and stored at −20 °C until assayed. Liquid chromatography-mass spectrometry was performed at the Brigham and Women’s Hospital Research Assay Core to determine concentrations of serum *β-*estradiol and progesterone.

#### DIVA

Endocrine measures include salivary [E_2_] and [P_4_] concentrations. Saliva samples were collected via passive drool into 2 mL sterile cryovials shortly after waking twice a week (3–4 days apart) and stored at −20 °C until shipping. Samples were shipped following completion of the study to Salimetrics’ SalivaLab (Carlsbad, CA). To assess [E_2_], samples were assayed using the Salimetrics Salivary Estradiol Assay Kit (Cat. No. 1-3702) with an assay range from 1 to 32 pg/mL, a lower sensitivity limit 0.1 pg/mL), and a 7.45% inter-assay coefficient of variation. To assess [P_4_], samples were assayed using the Salimetrics Salivary Progesterone Assay Kit (Cat. No. 1-1502) with an assay range from 10 to 2430 pg/mL, a lower sensitivity limit 5 pg/mL), and a 7.55% inter-assay coefficient of variation. Samples were assayed in duplicate without modifications to the manufacturers’ protocols. All samples had detectable levels of estradiol, but two samples from sub-03 had no detectable progesterone.

### Neuroimaging Data

#### 28andMe, 28andOC

MRI data were collected with a 3-Tesla Siemens Prisma scanner equipped with a 64-channel phased-array head coil at the University of California, Santa Barbara. Data include structural T1-weighted images and a 10-minute resting state scan, during which the participant was instructed to keep her eyes open. Structural T1-weighted images were acquired using a T1-weighted MP-RAGE sequence, with TR = 2500 ms, TE = 2.31 ms, TI = 934 ms, and 0.8 mm slice thickness. Resting-state functional images were collected using a multiband BOLD EPI sequence that acquired 72 oblique slices with TR = 720 ms, TE = 37 ms, a multiband factor of 8, interleaved acquisition, and a 52º flip angle.

#### DIVA

MRI data were collected with a 3-Tesla Siemens Prisma scanner with a 32-channel head/neck coil at Florida International University’s Center for Imaging Science (Miami, FL USA). Data used here, collected as part of a larger imaging battery, include structural T1-weighted images and two 5-minute resting-state scans, during which participants were instructed to remain still and keep their eyes on a fixation cross presented on a screen outside the bore. Structural T1-weighted images were acquired using a 3D T1w inversion prepared RF-spoiled gradient echo scan, the same sequence used by the Adolescent Brain and Cognitive Development (ABCD) Study℠ (Casey et al., 2018), with anterior-to-posterior phase encoding direction, TR = 2500 ms, TE = 2.88 ms, TI = 1070 ms, and 1 mm^3^ isotropic voxels. Resting state fMRI data were collected using a multiband, multi-echo (MBME) BOLD EPI sequence, from the distribution of multi-band accelerated EPI sequences (Moeller et al., 2010) developed by the Center for Magnetic Resonance Research at the University of Minnesota. The MBME GRE-EPI sequence acquired 48 slices with 2.4 mm^3^ isotropic voxels at a 30° transverse-to-coronal orientation with anterior-to-posterior phase encoding direction at each of 4 echoes (TE1 = 11.80 ms, TE2 = 28.04 ms, TE3 = 44.28 ms, TE4 = 60.52 ms) with TR = 1500 ms, a multiband acceleration factor of 3, interleaved acquisition, in-plane GRAPPA acceleration, a 77° flip angle.

### MRI Data Processing

Results included in this manuscript are derived from preprocessing performed using fMRIPrep 20.2.1 (Esteban, Markiewicz, et al. (2018); Esteban, Blair, et al. (2018); RRID:SCR_016216), which is based on Nipype 1.5.1 (Gorgolewski et al. (2011); Gorgolewski et al. (2018); RRID:SCR_002502). For more detail, see the supplementary methods.

#### Anatomical Image Preprocessing

For each participant, T1-weighted images were corrected for intensity non-uniformity using ANTS, then skull-stripped using OASIS30ANTs as a target template, and segmented into cerebrospinal fluid (CSF), white matter (WM), and gray matter (GM) using FSL’s fast. Nonlinear registration was then performed to align participants’ T1w images with ICBM 152 Nonlinear Asymmetrical template version 2009c (MNI152NLin2009cAsym).

#### Functional Image Processing

##### Single-Echo Image Processing: 28andMe, 28andOC

For each BOLD fMRI run, fieldmaps were estimated from EPI reference images with opposite phase encoding directions, and then used to create a corrected reference image for coregistration with the participant’s T1w image (6 degrees of freedom, using boundary-based registration). FSL’s MCFLIRT was used to estimate head motion parameters and slice-timing correction was performed using AFNI’s 3dTshift. Then, bias field correction and motion correction were applied in one step to resample each volume into the participant’s native space. These preprocessed data were then resampled into MNI152NLin2009cAsym space and confounding time series (i.e., of head motion, CSF, WM, GM, and their temporal derivatives) were extracted. Data were then high pass filtered and principal components of the CSF and WM voxels were extracted (i.e., per aCompCor). Finally, a motion outlier time series was created, flagging any volume with framewise displacement (FD) greater than 0.5 mm.

##### Multi-Echo Image Processing: DIVA

For each BOLD fMRI run, fieldmaps were estimated from EPI reference images with opposite phase encoding directions, and then used to create a corrected reference image for coregistration with the participant’s T1w image (6 degrees of freedom, using boundary-based registration). FSL’s MCFLIRT was used to estimate head motion parameters and slice-timing correction was performed using AFNI’s 3dTshift. Then, bias field correction and motion correction were applied in one step to resample each volume into the participant’s native space. From this data, a quantitative T2* image was estimated via a nonlinear fit to a monoexponential signal decay model using the maximal number of echoes with reliable signal for each voxel. This T2* image was then used to optimally combine the echoes at each TR to create the preprocessed BOLD data. These preprocessed data were then resampled into MNI152NLin2009cAsym space and confounding time series (i.e., of head motion, CSF, WM, and their temporal derivatives) were extracted. Finally, a motion outlier time series was created, considering any volume with framewise displacement (FD) greater than 0.5 mm.

#### Functional Connectivity Estimation

Following fMRI preprocessing, the Individual Differences in Connectivity pipeline (IDConn, v. 0.3; DOI: 10.5281/zenodo.6080388) was used to estimate connectivity per participant per session (Bottenhorn & Salo, 2024). First, Nilearn was used to mask preprocessed fMRI data with a 268-region atlas generated from via spectral clustering voxelwise resting-state functional connectivity data to define homogeneous, spatially-constrained clusters (i.e., regions), covering cortex, subcortex, and cerebellum (Craddock et al., 2012). After regressing out six motion parameters, CSF and WM signals, and their temporal derivatives, the average residual BOLD signal was extracted per region and standardized. BOLD signals were then correlated, region-wise, to assemble an adjacency matrix per participant per session. Thus, these adjacency matrices represent unthresholded functional connectome estimates as correlation-weighted connections (i.e., edges) between brain regions (i.e., nodes), pairwise, per time point per participant in each dataset.

Participants’ head motion was well within the acceptable range for resting-state fMRI data (i.e., mean framewise displacement <0.2 mm), indicating that no participant’s imaging data was sufficiently corrupted by motion. While head motion did differ between the two datasets, it was unrelated to HC use, [E_2_], or [P_4_] (Supplementary Table 1).

**Table 1.** Performance and generalizability of contraceptive- and hormone-related subnetworks.

|  | HC use | Estradiol |  | Progesterone |  |
| --- | --- | --- | --- | --- | --- |
|  |  | Base | HC use | Base | HC use |
| CV mean ± sd | 79.8%<br>± 11.4% | 0.130<br>± 0.277 | 0.025<br>± 0.304 | 0.357<br>± 0.252 | 0.189<br>± 0.296 |
| Alpha | $5.99 \times 10^{-3}$ | $7.74 \times 10^{-4}$ | $5.99 \times 10^{-3}$ | $1.00 \times 10^{-4}$ | $5.99 \times 10^{-3}$ |
| In-sample (28and*) |  |  |  |  |  |
| Model performance | 74% | 0.937 | 0.814 | 0.804 | 0.950 |
| MSE | 0.0833 | 0.174 | 0.368 | 0.0861 | 0.0172 |
| Out-of-sample (DIVA) |  |  |  |  |  |
| Model performance | 50% | 0.182 | 0.648 | 0.429 | -0.357 |
| MSE | 0.5 | 0.619 | 0.0810 | 0.0719 | 0.118 |

*Note.* HC use models are logistic regression with $L_2$ penalties and their performance is percent accuracy of HC use predicted by functional connectivity. Estradiol and progesterone models are linear regression with $L_2$ penalties (i.e., Ridge regression) and their performance is provided by the Spearman correlation between true hormone concentrations and hormone concentrations predicted from functional connectivity of the identified subnetwork. Results from sensitivity analyses are shown in Supplemental Table

### Inferential Statistics

Data used to conduct these analyses can be found at github.com/62442katieb/menstrual_cycle_connectomics.

#### Missing Data Across Variables

First, we assessed missingness per variable by calculating (1) the proportion of missing values per participant per variable and (2) point-biserial correlation coefficients between missingness on each variable and the values of each other variable, to determine whether variables were missing at random or not.

Missing data was <10% per participant per variable and there was no evidence to suggest missingness on any one variable was conditioned on values of other variables, providing no sufficient evidence that missingness is not completely at random.

#### Analysis 1: Connectome Similarity

The degree of similarity in functional connectivity across the whole brain, i.e., connectome similarity, due to each (1) group (i.e., shared across participants), (2) dataset, (3) individual, and (4) hormonal contraceptive use was quantified by computing a product-moment correlation coefficient for each pair of networks (i.e., the upper triangle of the adjacency matrix). Then, whole-brain network similarity was calculated as the average of Fisher-transformed product-moment correlations for (1) all individuals (i.e., *group*-level similarity) (2) each *dataset*, (3) *HC use* status and (4) each *individual*, respectively. The whole-brain network similarity values were then compared between each of the subdivisions using two-sample *t*-tests, evaluated at a significance threshold of *α* < 0.01.

#### Analysis 2: Hormone-Related Connectomics

Finally, we used a predictive modeling approach to identify intrinsic functional subnetworks related to HC use and within-individual differences in hormone levels across the menstrual and HC pill cycles, and then applied transfer learning to test their generalizability out-of-sample (i.e., between individuals).

First, a Python adaptation of NBS-Predict (Serin et al., 2021), included in IDConn (Supplementary Figure 1), was used to identify functional connectivity related to HC use in a 5-fold cross-validation framework with hyperparameter tuning to determine the optimal *L*_2_ coefficient (i.e., *α*). To do so, data from 80% of the time points (i.e., 48 sessions) in 28andMe and 28andOC were used to train a model, following within-fold regression of average head motion from each edge, by (1) performing connection-wise *F*-tests for connectivity differences between naturally cycling (NC) and HC-using timepoints, and (2) selecting functional connections that were significantly associated with the HC use above a given *p*-threshold (i.e., *p*(*F*) < 0.05). From these suprathreshold connections, (3) NBS-Predict determined the HC-associated subnetwork, ***N***_***HC***_, as the largest connected component (i.e., subnetwork of suprathreshold connections in which each brain region is connected to every other brain region either directly or indirectly), removing the connections that were not part of that component. The strengths of these connections were, then, used as features in a logistic regression model of the form in Equation 2, regressing HC use onto subnetwork connectivity in the remaining 20% of time points (i.e., 12 sessions), to estimate how related the subnetwork as a whole is to HC use.

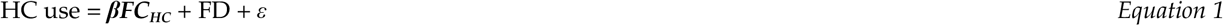

This process was repeated 1000 times, randomly sampling the training dataset (28andMe + 28andOC) to yield a distribution of HC-related subnetworks and their accuracy in classifying HC use. Then, an average of these subnetworks, weighted by their classification accuracy, was calculated and thresholded to determine the functional subnetwork most associated with HC use. Finally, the connectivity of this subnetwork was used to train a logistic regression model on the full training dataset (i.e., all 60 sessions from 28andMe + 28andOC) and the resulting model parameters (i.e., *α, β*) were used to predict HC use from connectivity of the subnetwork in DIVA data and estimate generalizability from out-of-sample predictive accuracy.

Second, NBS-Predict was similarly used to separately predict [E_2_] and [P_4_] from functional connectivity in a 5-fold cross-validation framework, by (0) regressing HC use and framewise displacement from [E_2_], [P_4_], and each edge, (1) calculating connection-wise *F*-tests between the residualized connectivity and [E_2_] and [P_4_] in 80% of 28andMe and 28andOC time points (i.e., hormone-related connectivity accounting for HC use) to (2) select suprathreshold connections (i.e., *p*(*F*) < 0.05) and (3) assemble separate estradiol- and progesterone-associated subnetworks, *N*_*E2*_ and *N*_*P4*_, from the largest connected components. Then, linear regression with *L*_2_ regularization (i.e., ridge regression) was used to predict the hormone levels, per Equation 3, in the remaining 20% of 28andMe and 28andOC data. Model performance was assessed via correlations between actual and predicted hormone concentrations, per NBS-Predict methodology (Serin et al., 2021), but using Spearman’s correlation instead of product-moment correlations to accommodate unmet distributional assumptions. Each model was also run without regressing out HC use (see parentheticals in *Equations 2-3*), to examine the role of exogenous hormones on connectivity related to endogenous hormones, in this context.

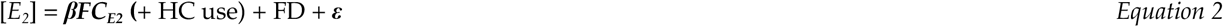

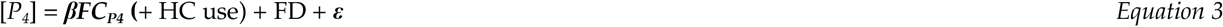

As above, this process was repeated 1,000 times and the resulting subnetworks and Spearman correlations were used to assemble weighted average estradiol- and progesterone-associated subnetworks, weighted by model performance. These subnetworks were then thresholded, retaining 50% of the most positively outcome-related edges, and applied to the full training dataset to train linear regression models for each [E_2_] and [P_4_]. The resulting model parameters (i.e., *α, β*) were then used to predict [E_2_] and [P_4_] from DIVA subnetwork connectivity data, to determine their out-of-sample generalizability.

Finally, we ran *post hoc* sensitivity analyses, dropping the high [E_2_] days (i.e., ovulatory peaks, menstrual cycle days 12-16) from the data used identify HC- and hormone-related subnetworks (i.e., training dataset: 28andMe, 28andOC, as those days were not sampled in the DIVA dataset and brain-hormone associations vary across menstrual cycle phases (Casto et al., 2022).

## Results

Hormone concentrations across the measurement periods (scaled between 0 and 1, within each dataset, for comparison across measurement techniques) are displayed in Figure 3, showing differences in cycle timing and measurement period across subjects. Here, “cycle day” refers to the number of days since the start of the participant’s most recent menses.

**Figure 2.**
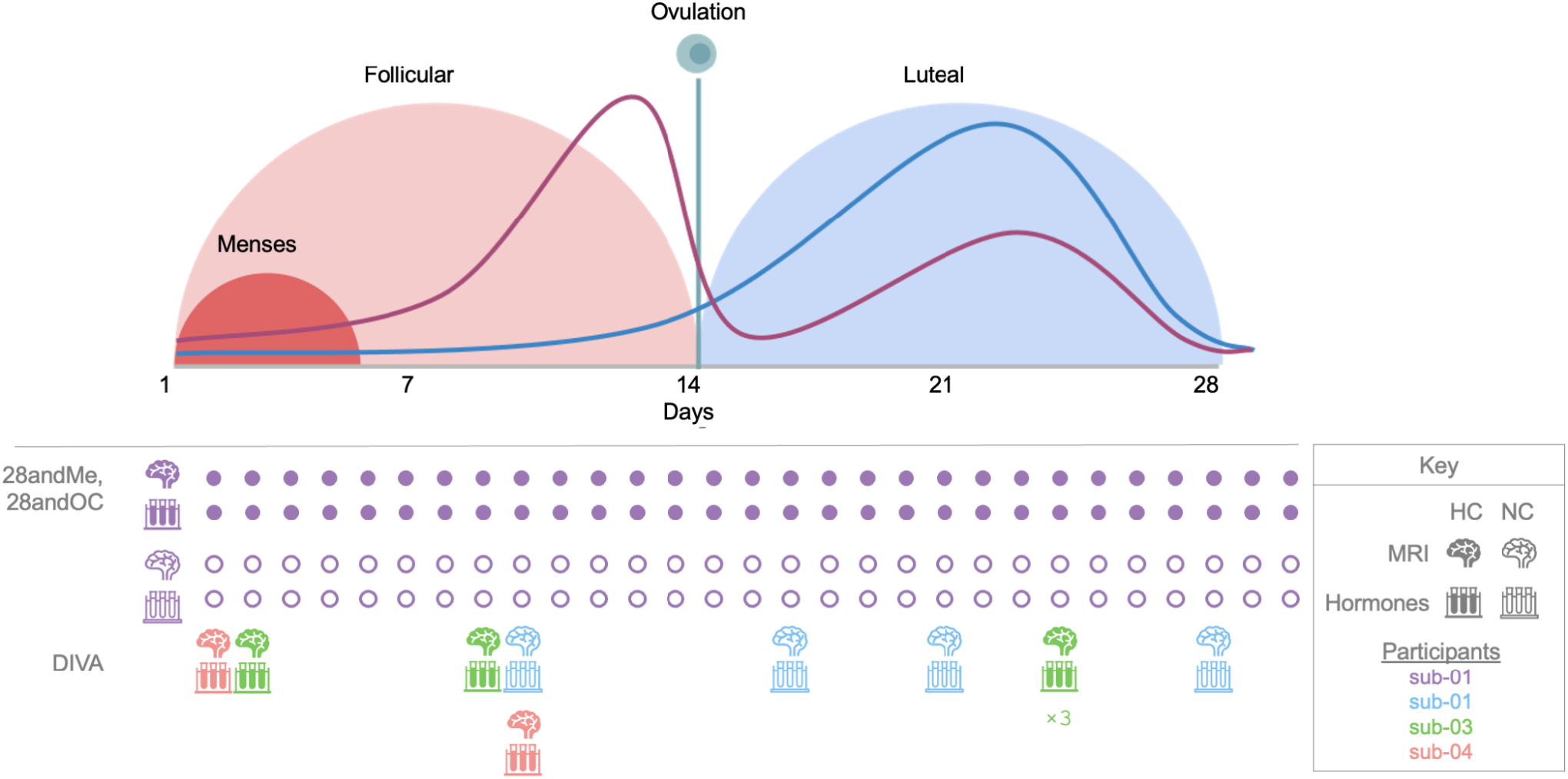
Schematic showing the timing of endocrine and neuroimaging data collection per dataset. The graph (top) is a schematic illustration (i.e., not derived from data presented here) of the phases of the ovarian cycle and standard hormone concentrations throughout. The purple line represents estrogen levels, while the blue line represents progesterone levels (not to scale between hormones). This schematic is based on a 28-day menstrual cycle, which is the average length but not representative of all women. The follicular phase length varies more between individuals, while the luteal phase length is generally more consistent. The timing of data collection for each dataset is presented below. Brain icons indicate days on which MRI data were acquired, test tube icons indicate biospecimen collection days, both of which are color-coded by participant. Filled icons indicate data collected from a participant using HCs, unfilled icons indicate data collected from a naturally cycling participant. After the first data points from 28andMe and 28andOC, subsequent data collection days are abbreviated with filled and unfilled circles, respectively, to more simply indicate data collection scheme.

**Figure 3.**
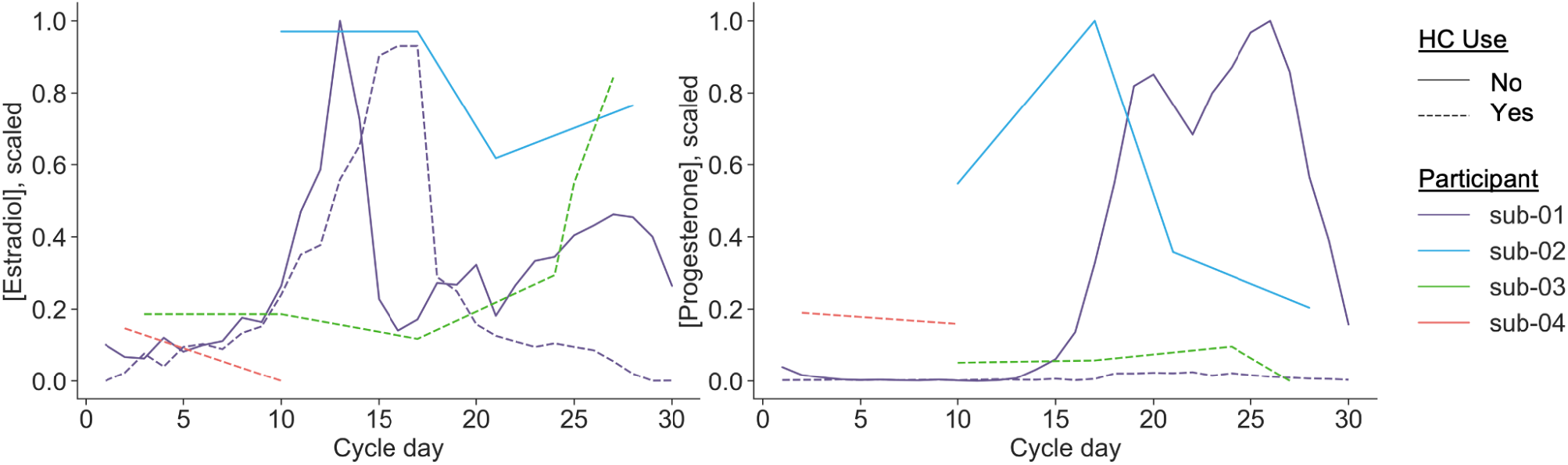
Dataset-scaled estradiol and progesterone concentrations across the sample. Estradiol (left) and progesterone (right) were assessed via assay of either saliva or blood samples, as the collection method differed between datasets. Shown here are estradiol and progesterone concentrations per participant across the menstrual cycle, scaled between 0 and 1 within each dataset (i.e., 28andMe/28andOC; DIVA). Solid lines indicate freely cycling participants (i.e., sub-01, sub-03), while dashed lines indicate participants using hormonal contraceptives (i.e., sub-01, sub-02, sub-04). Variations in length and reporting of menstrual cycle cause lags between estradiol and progesterone increases seen in menstruating women. Error bar on sub-04, cycle day 24 indicates multiple measurements on that day, although this is likely due to a reporting error. Thus, cycle day is not used in any of the following analyses.

### Analysis 1: Connectome Similarity

Overall, networks were more similar within individuals than between individuals (Figure 4A). In line with results from Gratton et al. (2018), we found that network similarity (Figure 4B) within each individual was more than twice the magnitude of similarity between individuals (i.e., across the group). Additionally, we found that network similarity was of similar magnitude between dataset and HC use status, though group-level similarity was significantly greater than similarity based on HC use status (*p* < 0.01). However, as one dataset (i.e., 28andMe, 28andOC) was limited to one individual, these results should be interpreted with caution and warrant further investigation in a larger multi-site dataset.

**Figure 4.**
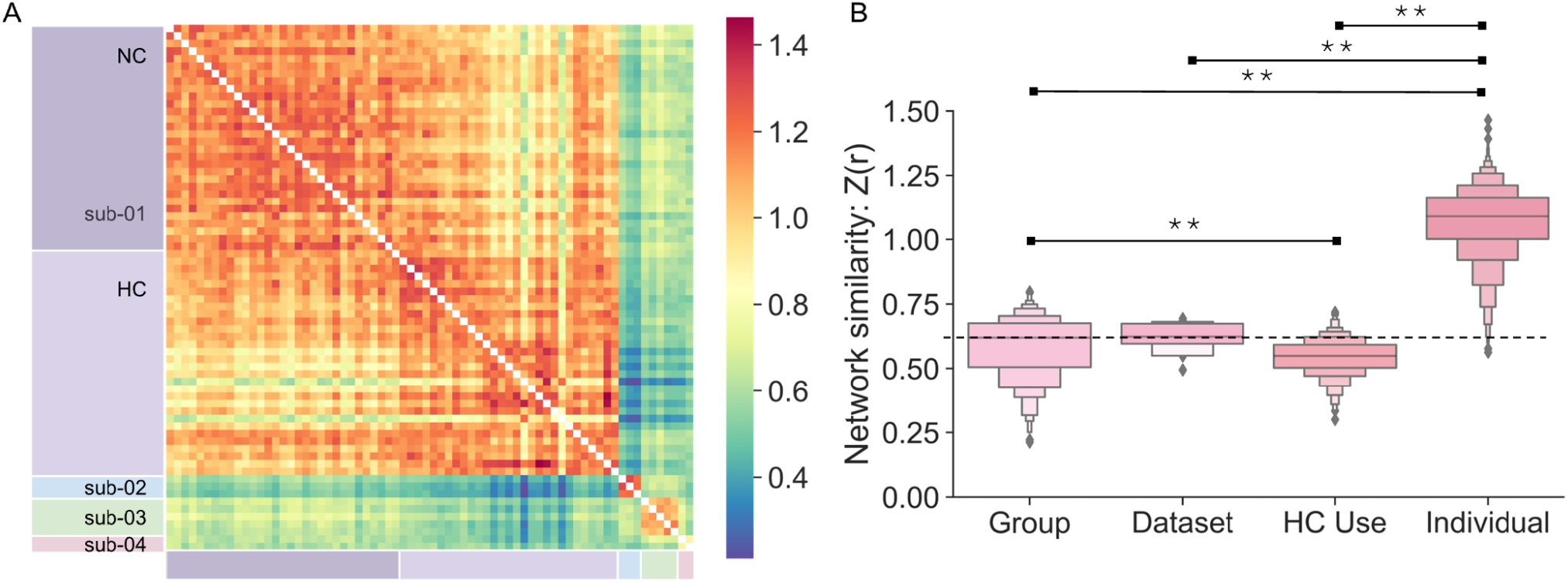
Network similarity across participants and sessions. A: Network similarity is shown as correlation coefficients Fisher-transformed for normality (*Z*(*r*)). Rows/columns are sorted by participant and session. B: Network similarity is calculated across the group (from different individuals, datasets, HC use strategies), between datasets (from different individuals and HC use strategies), across HC use strategies (from different individuals and datasets), and between individuals (from different datasets and HC use statuses). Asterisks indicate significant differences in network similarity, *p* <0.01. Abbreviations: NC: naturally cycling, HC: hormonal contraceptives.

### Analysis 2: Contraceptive- and Hormone-Related Connectomics

NBS-Predict revealed diffuse, brain-wide subnetworks associated with HC use, as well as more limited [E_2_]- and [P_4_]-related subnetworks, with varying generalizability to independent data (Table 1).

#### Contraceptive-related functional connectivity changes

Within-individual contraceptive-related functional connectivity was robust and included nearly every region of the brain (Figure 5). Average model accuracy across model-building iterations (i.e., assessed on an unseen subset of 28andMe, 28andOC data) was 79.8% and overall in-sample accuracy was 74%. When generalizability of the contraceptive-related subnetwork was assessed on an independent sample (i.e., DIVA), accuracy was 50%. There was little variation, spatially or between model-building iterations, in the predictive power of functional connections across the brain (Figure 5A,B). Regionally, the insula, caudate, midcingulate cortex, superior postcentral gyri, and orbitofrontal cortex displayed the greatest contraceptive-related connectivity, while the cerebellum and medial temporal lobe displayed the least. Across large-scale brain networks, somatomotor and ventral attention networks displayed the greatest contraceptive-related connectivity, while cerebellar, limbic, and visual networks displayed the least (Supplementary Figure 2, top row).

**Figure 5.**
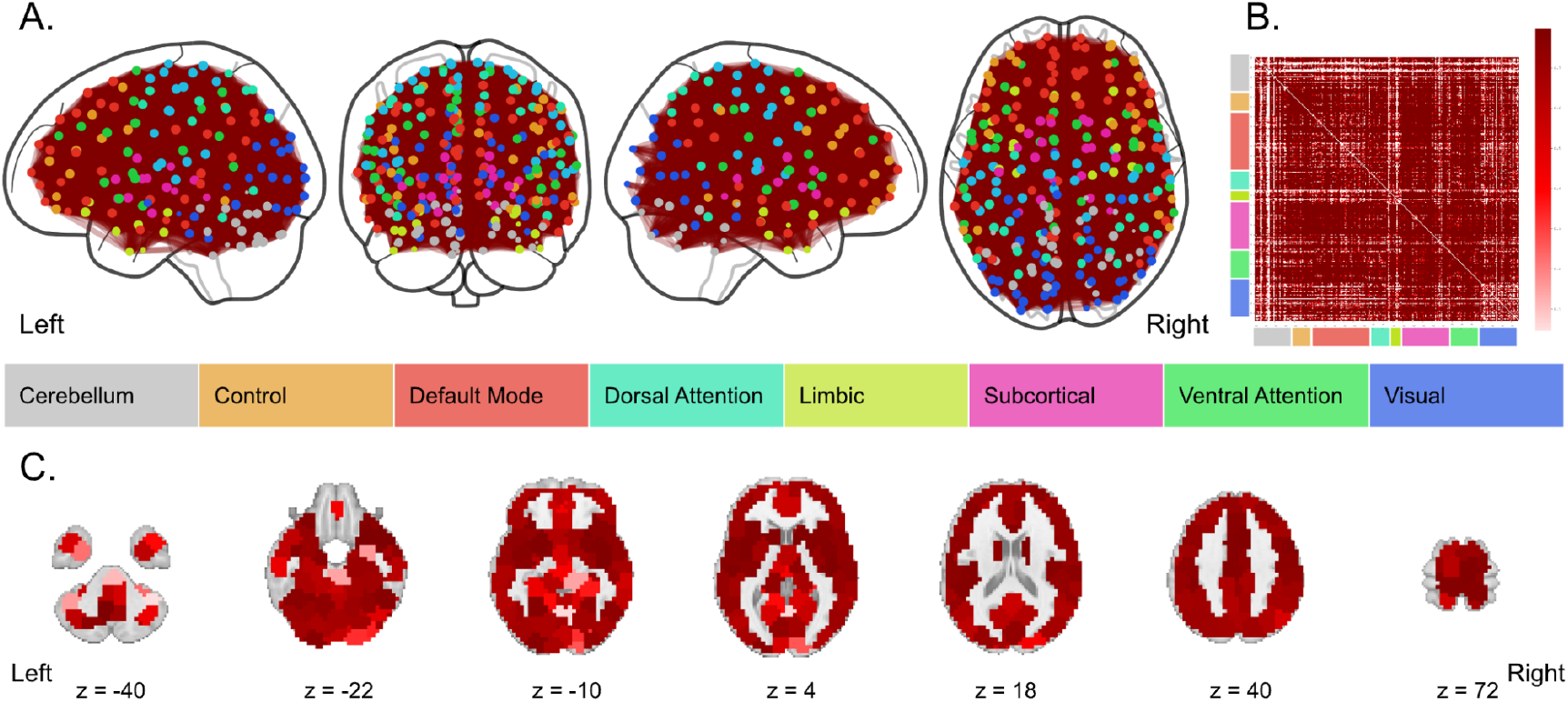
Functional connectivity predictive of HC use. A. The top 0.001% of functional connections most predictive of HC use across training folds. Node color is determined by the functional network, as defined by Yeo et al. (Yeo et al., 2011) with which each region demonstrates the greatest overlap. Edge color is determined by the average accuracy of the identified subnetwork in predicting HC use in the held out sample, across training folds. Average accuracy was consistently high, that most edges shared the same, maximum value (i.e., 0.799) B. Average accuracy of all functional connections in predicting HC use across model training. C. HC-use-associated node strength, or the sum of each brain region’s accuracy-weighted functional connections, with *z*-coordinates of each slice given in MNI space..

#### Hormone-related functional connectivity

Estradiol-related functional connectivity was sparser (Figure 6; Supplementary Figure 1, middle row). Average Spearman correlation between actual and predicted estradiol concentrations, within 28andMe, 28andOC data, was 0.130 (± 0.277) without HC use as a covariate and 0.025 (± 0.304) with HC use as a covariate. Thus, accounting for HC use caused the identified subnetworks to more accurately describe [E_2_] in the held-out sessions, but with a wider spread, potentially indicating more variability in the edges identified as related to [E_2_]. Once the performance-weighted subnetworks were thresholded to only include the top 50% of [E_2_]-related connections (i.e., with average *r* > 0 across training folds), overall in-sample correlation was 0.168 without HC use as a covariate and 0.0392 with HC use as a covariate. When the [E_2_]-relatedness of the same connectivity was assessed in an independent dataset (i.e., DIVA Study data), out-of-sample correlation was −0.491 without HC use as a covariate and 0.648 with HC use as a covariate. Thus, the identified functional subnetwork demonstrated moderate performance in estimating [E_2_] within an individual, its generalizability to other individuals depended on HC use. When accounting for HC use, the identified subnetwork performed better within an individual, but worse in independent data. Neuroanatomically, the right anterior insula demonstrated the highest node strength in the [E_2_]-related subnetwork (Figure 6A, C). Accounting for HC use, estradiol-related connectivity was greatest in ventral attention network (VAN) regions, most notably in the anterior insula and between VAN and somatomotor regions. In contrast, connectivity of virtually no dorsal attention, limbic, or subcortical regions were related to changing [E_2_] (Supplementary Figure 1, middle row).

**Figure 6.**
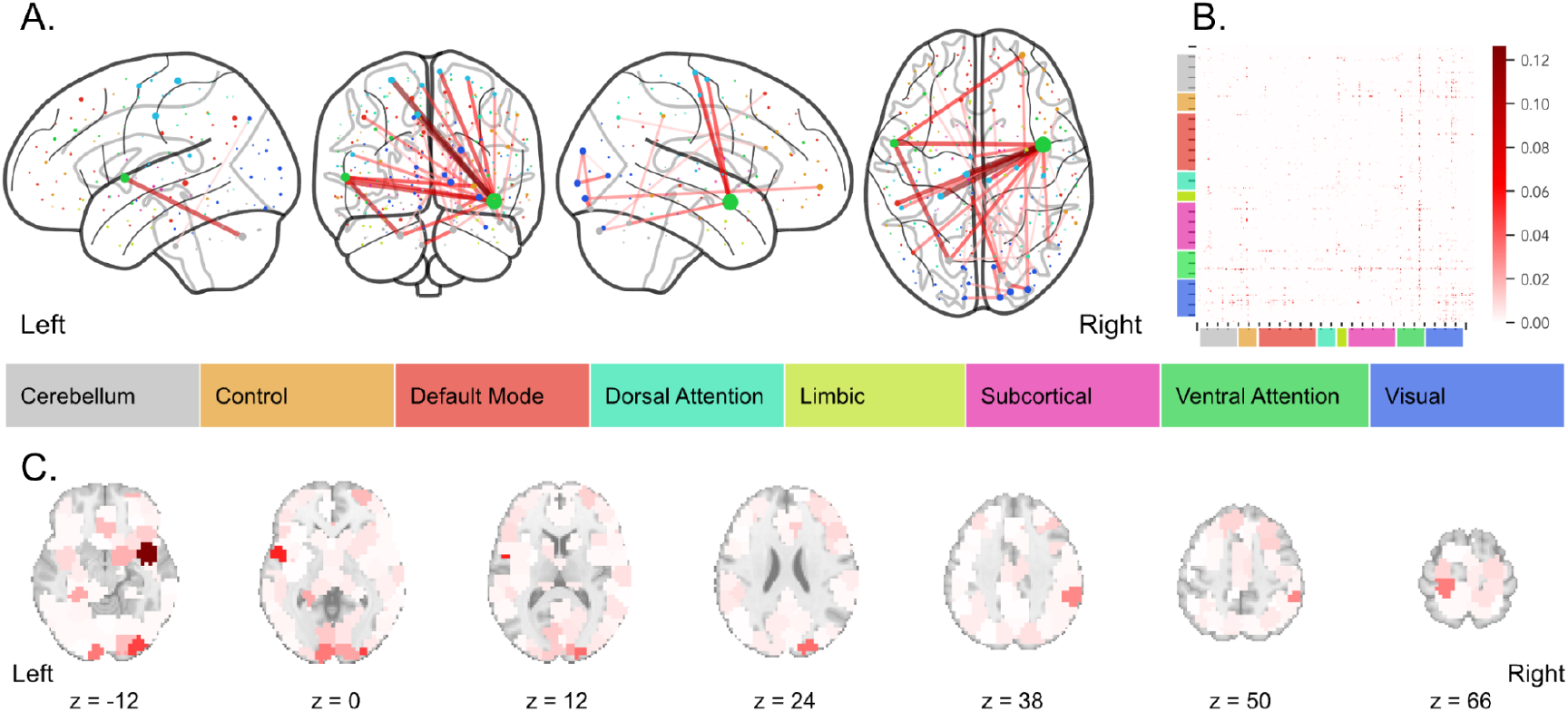
Functional connectivity predictive of estradiol levels. A: The top 0.15% of functional connections most predictive of [E_2_] across training folds. Node color is determined by the functional network, as defined by Yeo et al. (Yeo et al., 2011) with which each region demonstrates the greatest overlap. Edge color is determined by the average Spearman correlation (i.e., *r*) of predicted [E_2_] vs. actual [E_2_] across training folds. B: Weighted average subnetwork, using Spearman correlation (i.e., *r*) of model’s predicted [E_2_] vs. actual [E_2_] across training folds. C: [E_2_]-associated node strength, or the sum of each brain region’s *r*-weighted functional connections, with *z*-coordinates of each slice given in MNI space..

Compared with estradiol, progesterone-related functional connectivity was considerably denser and more widespread (Figure 7). The final model’s Spearman correlation between actual and predicted serum progesterone concentrations (i.e., tested on an unseen subset of 28andMe, 28andOC data), was 0.80 (*p* < 0.01; MSE: 0.086) without HC use as a covariate and 0.95 (*p* < 0.01; MSE: 0.0172) with HC use as a covariate. Thus, accounting for HC use caused the identified subnetworks to more accurately reflect changing [P_4_], potentially due to a confounding blunting of progesterone levels by HC use. Throughout the network-identification process, adding HC use yielded poorer performance and a greater range of model fit, indicating greater variability in the identified networks. The final subnetwork, however, was a more accurate reflection of progesterone-related functional connectivity in that individual when accounting for HC use, but showed less generalizability to other individuals, as shown by greater error in [P_4_] estimated from others’ functional connectivity (without HC: *r* = 0.49, MSE: 0.0719; with HC: *r* = −0.357, MSE = 0.118). Unlike with [E_2_]-related functional subnetworks, adjusting for HC use caused the identified functional subnetwork’s estimation of [P_4_] in novel data to perform worse than did the identified functional subnetwork that did not adjust for HC use. Accounting for HC use, the progesterone-related subnetwork demonstrated the most connectivity between cerebellar and ventral attention regions, within the visual network, and between limbic regions and each subcortical, dorsal attention, and visual regions. In contrast, other visual connectivity, frontoparietal control network connectivity, and default mode network connectivity was less related to changing [P_4_] over the menstrual cycle.

**Figure 7.**
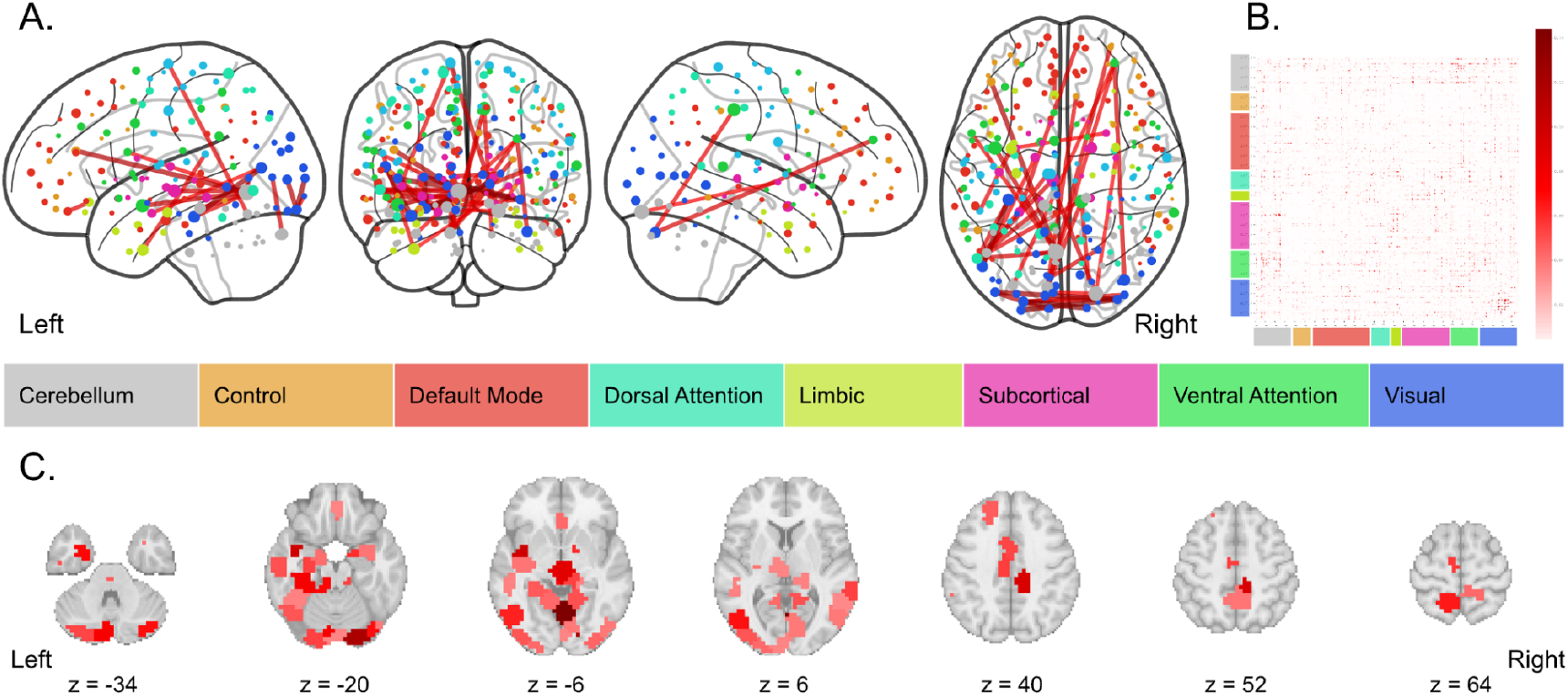
Functional connectivity predictive of progesterone levels. A: The top 0.15% of functional connections most predictive of [P_4_], across training folds, accounting for HC use. Node color is determined by the functional network, as defined by Yeo et al. (Yeo et al., 2011) with which each region demonstrates the greatest overlap. Edge color is determined by the average Spearman correlation (i.e., *r*) of predicted [P_4_] vs. actual [P_4_] across training folds. B: Weighted average subnetwork, using Spearman correlation (i.e., *r*) of model’s predicted [P_4_] vs. actual [P_4_] across training folds. C: [P_4_]-associated node strength, or the sum of each brain region’s *r*-weighted functional connections, with *z*-coordinates of each slice given in MNI space.

#### Sensitivity analyses: Removing ovulatory peaks from training dataset

We repeated all predictive connectomics analyses after excluding time points from the training dataset that fell within the ovulatory window (Supplementary Table 2). Without the high-estradiol days within the ovulatory window, the HC-related subnetwork exhibited poorer in-sample model fit and worse generalizability to DIVA Study data (i.e., 50% accuracy). Compared with the models presented above, the [E_2_]-related subnetwork trained without ovulatory window data exhibited better in-sample model fit when accounting for HC use, worse in-sample model fit when not accounting for HC use, and poorer generalizability in both cases. Compared with the models presented above, the [P_4_]-related subnetwork trained without ovulatory window data, on the other hand, demonstrated worse in-sample performance when accounting for HC use, but better in-sample model fit when not accounting for HC use, and better generalizability to DIVA data regardless of accounting for HC use. The topography of the HC-related subnetwork identified without ovulatory window data was virtually identical to the subnetwork presented above: widespread, highly-related connectivity. The topography of the [E_2_] and [P_4_]-related subnetworks (i.e., with and without accounting for HC use) were largely unchanged following the removal of ovulatory window data (Supplementary Figure 3).

## Discussion

Steroid hormones are critical neuromodulators. As human neuroimaging strives toward population neuroscience and precision medicine, understanding hormones’ roles in brain function, as well as idiosyncrasies and commonalities therein, becomes paramount. First, these results replicate prior findings from Gratton and colleagues: stable features common across the group and specific to individuals contribute almost equally to functional connectome similarity (Gratton et al., 2018). Further, we extend this work to show that individual-specific features continue to contribute significantly to connectome similarity, beyond differences in datasets and participant HC use. Second, predictive modeling identified widespread functional connectivity changes associated with HC use, and sparser changes related to estradiol and progesterone fluctuations throughout the menstrual cycle. The identified HC- and hormone-related connectivity exhibited mixed generalizability to other individuals, potentially reflecting differential brain–hormone associations depending on HC progestin formulations. Altogether, this work marks an important step forward in understanding neuroendocrine dynamics inherent to the adult brain in people who menstruate and begs further study of contraceptive-related differences and functional connectomics across the menstrual cycle.

### Individual-specific functional connectivity

The brain’s functional network organization is relatively idiosyncratic from birth (Wang et al., 2021) into adulthood (Gratton et al., 2018) in both its neuroanatomy (Seitzman et al. 2019) and connectomics (Finn et al., 2015). Here, we replicate prior findings regarding approximately equal influences of group commonalities and individual-specific factors on functional connectomics (Gratton et al., 2018). Using data from two independent datasets with adult female participants of similar age (i.e., 23 to 31 years), we found that the magnitude of contributions of dataset differences and imaging sequence (inextricable in these data) to network similarity are not significantly different than that of group commonalities, although HC-use contributes slightly, but significantly less, than do group commonalities. Even in the face of a common exogenous hormone manipulation from HC use, individual traits dominate functional connectivity. The relative contributions of HC use and individual factors to functional connectivity are particularly interesting on two fronts. First, there is evidence that the menstrual cycle varies as much within individuals as between, which neatly aligns with our findings. That is, after accounting for similarity due to shared group factors (i.e., between-individual similarity; z = 0.56), the magnitude of additional similarity due to individual factors (i.e., z = 1.04 − 0.56 = 0.48) is similar. Second, the observed ubiquity of within-individual HC-related functional connectivity changes, in contrast with near-chance accuracy in predicting HC use in other individuals, reflects the contribution of shared-group vs. individual-specific factors contributing to functional connectivity. That is, a similar exogenous hormone manipulation is robustly related to brain-wide changes in connectivity, but is not more distinctive than other individual differences in connectivity.

### Contraceptive-related changes in functional connectivity are ubiquitous

Hormonal contraceptive use impacted functional connectivity of every large-scale functional brain network, with the smallest impacts in cerebellar, visual, and limbic regions and the largest impacts in dorsal and ventral attention networks, as well as frontoparietal control, somatomotor, and default mode networks. As these subnetworks were identified using data from only one individual, their generalizability to other individuals provides additional information about the extent to which individual-specific traits inform neuroendocrine fluctuations during the adult menstrual cycle and the extent to which hormonal contraceptives affect these fluctuations. Two notable findings arose from assessing generalizability of the identified endocrine-related subnetworks. First, accounting for HC use in models identifying hormone-related connectivity improved our ability to identify progesterone-related connectivity but worsened our ability to identify estradiol-related connectivity. Second, and conversely, accounting for HC use in uncovering hormone-related connectivity improved the generalizability of estradiol-related connectivity, but substantially worsened the generalizability of progesterone-related connectivity. Although these results represent many observations from only a few individuals, they recapitulate prior findings regarding the extent to which HC use alters functional connectivity, including diffuse, altered connectivity between several large-scale brain networks (Casto et al., 2022; Petersen et al., 2014).

Differences in model fit due to accounting for HC use indicate that estradiol- and progesterone-related connectivity is differentially confounded by the exogenous hormones in oral contraceptives. However, accounting for HC use yielded an estradiol-related subnetwork *more* robust to individual differences, but rendered the progesterone-related subnetwork *less* robust. The participants in these studies all used HCs with different progestin formulations and different amounts of ethinyl estradiol, which makes it difficult to disentangle individual-specific and formulation-related differences. The diminished generalizability of progesterone-related functional connectivity may reflect differential effects on functional connectivity between progestins. This possibility is bolstered by the fact that the progestins in each HC formulation used by participants in this study have different pharmacological and neurobiological effects (Bullock & Bardin, 1977; Casto et al., 2022; Griksiene et al., 2022). In addition to their progestational effects, norethindrone acetate (used by sub-03 in DIVA) has strong androgenic but weak estrogenic effects, norgestimate (used by sub-04 in DIVA) has androgenic effects and antiestrogenic effects, and levonorgestrel (i.e., used by sub-01 in 28andOC data) has high androgenic effects, but weak estrogenic effects (Griksiene et al., 2022). Further research including a larger sample size with measurements of synthetic hormones (e.g., ethinyl estradiol, levonorgestrel) in multiple, independent datasets would be required to support this theory and clarify the interactions between exogenous, synthetic hormones from HCs and endogenous hormones.

### Hormone-related functional connectivity changes across the menstrual cycle

We identified estradiol- and progesterone-related subnetworks that exhibit changing functional connectivity across the menstrual cycle, using two months of daily biospecimen and functional neuroimaging data. The estradiol-sensitive subnetwork was dominated by ventral attention, visual, and cerebellar network regions with a the core subnetwork structure (i.e., bilateral anterior insula, occipital, orbitofrontal, (para)hippocampal, and cerebellar connectivity) that persisted when accounting for HC use. The progesterone-sensitive subnetwork, however, was more widespread. Accounting for HC use diminished progesterone-related connectivity affecting its core subnetwork structure (i.e., diminishing connectivity of midline frontal and precuneus regions while increasing within-occipital lobe connectivity). Robustly identifying progesterone-related functional connectivity is further complicated by unmeasured levels of progesterone synthesized in the brain that are distinct from circulating progesterone (measured here) and modulated by estradiol (Micevych & Sinchak, 2008).

Much of the extant literature describing brain changes along the menstrual cycle lack methodological rigor and statistical power (Dubol et al., 2021), making it difficult to reliably contextualize these findings, as does this study’s small sample and demonstrated connectome idiosyncrasy. Some have identified no effects of menstrual cycle fluctuations on resting-state functional connectivity (De Bondt et al., 2015; Hjelmervik et al., 2014), though this may be due to insufficient statistical power with 16–18 naturally cycling female participants and only 3 time points each (i.e., once per menstrual cycle phase). However, a few consistent findings align with our results. A recent dynamic causal modeling study scanned 60 naturally cycling women 3 times (i.e., in early follicular, pre-ovulatory and mid-luteal phases) and uncovered estradiol-related insula connectivity with hemispheric differences, as seen here (Hidalgo-Lopez et al., 2021). A recent meta-analysis identified convergent gray matter volume and task-based activation differences in the insula over the course of the menstrual cycle (Dubol et al., 2021). Prior functional connectivity studies have also identified estradiol-related connectivity of occipital, inferior frontal, and temporal cortex regions (Craig et al., 2008; Engman et al., 2016; Meeker et al., 2020). Interestingly, after accounting for HC use in the current study the estradiol-related functional subnetwork involved relatively little limbic and subcortical connectivity, despite their relatively high estrogen receptor expression (i.e., ERα and, to a lesser extent, ERβ; (Österlund et al., 1999, 2000)) and estradiol concentrations (Bixo et al., 1995). This may be due to shared variance in subcortical–limbic connectivity between exogenous and endogenous estradiol levels, but it is unlikely given that subcortical and limbic regions play a relatively small role in the identified HC-related subnetwork. Thus, there may be interaction effects between estradiol and HC actions in subcortical and limbic connectivity that warrant future study. Finally, our results did not include the expected estradiol-related prefrontal cortex (PFC) connectivity. Instead, we found a minimal role for PFC connectivity in the estradiol-related subnetworks, regardless of accounting for HC use. Prior research indicates that PFC connectivity changes between the luteal and follicular phases of the menstrual cycle (Engman et al., 2018; Meeker et al., 2020; Wetherill et al., 2016), which may explain why it was not revealed here, as we did not incorporate menstrual cycle phase in these models.

### Limitations and Future Directions

Incorporating multiple datasets with different sampling rates and data collection methods has both strengths and limitations. Data in 28andMe and 28andOC was collected from a single participant, separated by one year with a 10-month HC wash-in period. Thus, in training the models, elapsed time, and HC use cannot be entirely disambiguated. Thus, the observed HC-related functional connectivity may also be related to differences in brain function seen over a year. This effect of time likely contributes to the poor generalizability of the identified HC-related subnetwork and is at least partially accounted for in the hormone models that account for HC use. The other consequence of study design is that these data cannot be used to address lagged associations between hormone concentrations and functional connectivity (as previously demonstrated by Pritschet and colleagues (Pritschet et al., 2020)), as DIVA Study participants only have biweekly, salivary hormone estimates and weekly MRI data. Furthermore, these results only represent a small proportion of potential HC-related functional connectivity and HC-related alterations of hormone-related functional connectivity, as only three HC formulations are represented in these data with one formulation per participant. Future research should include and compare oral contraceptive formulations (e.g., androgenic vs. antiandrogenic progestins, progestin-only mini pill), as well as other forms of hormonal contraceptives (e.g., implant, shot, intrauterine devices), while measuring concentrations of both exogenous and endogenous hormones.

In contrast to other multi-dataset investigations, we did not perform neuroimaging data harmonization between 28andMe, 28andOC, and DIVA Study data. In using independent datasets for model development and validation, we chose not to harmonize the neuroimaging data to avoid data leakage and any resulting bias of model coefficients or performance (Rosenblatt et al., 2024). Thus, our results regarding out-of-sample model performance reflect the generalizability of the identified contraceptive- and hormone-related subnetworks *despite* differences in MRI sequences and scanners, as well as differences in hormone estimates (i.e., salivary vs. serum), experimental design, and other confounding factors (e.g. time of day, temperature, air quality, season). While salivary and serum estimates of progesterone are generally highly correlated (Ellison, 1993), associations between salivary and serum estimates of estradiol can vary between individuals, potentially due to differences in salivary gland function (Lu et al., 1999) and procedures for estimating estradiol from saliva (e.g., immunoassays) are less reliable than those used to estimate estradiol from serum samples (e.g., liquid chromatography mass spectrometry) (Newman & Handelsman, 2014). Generalizability of estradiol-related connectivity identified here may be influenced by these differences in hormone estimation.

Finally, these analyses do not address the potential of a changing brain–hormone relationship across the menstrual cycle, as has been identified previously (Syan et al., 2017). The limited data available are insufficient to address different brain–hormone associations between menstrual cycle phases. Interestingly, 28andMe and 28andOC included brain and hormone measures collected during the high-estradiol ovulatory window, but the DIVA Study did not, and when ovulatory window data was removed in sensitivity analyses, both [E_2_]- and [P_4_]-related networks were topographically unchanged but [E_2_]-related subnetworks showed poorer generalizability, while [P_4_]-related subnetworks showed greater generalizability. Prior work with the 28andMe data showed that high-estradiol days during the ovulatory window were driving their estradiol-related connectivity findings (Pritschet et al., 2020). The marginally worse generalizability of these models in a dataset that did not measure the high-estradiol ovulatory window is curious. Future work with larger samples, and more within-individual data across menstrual cycle phases than is available in the DIVA Study, will help better understand the impact of cycle phase and ovulatory estradiol peaks on hormone-related connectivity.

## Conclusions

Here, we used two densely-sampled longitudinal datasets to assess individual-specific features of the functional connectome; to identify contraceptive-, estradiol-, and progesterone-related functional subnetworks across the menstrual cycle; and to assess the generalizability of those subnetworks. Our findings corroborated prior work regarding the balance of common and individual-specific contributions to functional brain organization, while extending the literature to consider the contribution of HC use. Further, we identified brain-wide changes in functional connectivity associated with HC use, which may be more idiosyncratic than they are generalizable, and smaller hormone-related subnetworks. Interestingly, accounting for HC use improved the generalizability, but not accuracy, of estradiol-related connectivity, but had the opposite effect on progesterone-related connectivity. Altogether, these findings hint at individual and progestin-related differences in functional brain network organization across the menstrual cycle, but underscore the need for larger studies of contraceptive-related changes in brain function with sufficient statistical power to compare potential effects of different HC formulations As the study of human neuroendocrinology advances with dense-sampling and precision approaches, it is imperative that women not be left behind.

## Acknowledgments

A special thank you to the individuals who participated in these studies.

Research described in this article was supported by NIH U01-DA041156 (ARL, KLB), R01-ES032295 (MMH, KLB), R01-ES031074 (MMH, KLB), and K99-MH135075 (KLB). Additional support was provided by the Office of Research and Economic Development (ORED) and the Dissertation Year Fellowship (DYF) Program at Florida International University (FIU). Finally, the authors would like to thank the FIU Instructional & Research Computing Center (IRCC, http://ircc.fiu.edu) for providing the HPC and computing resources that contributed to the research results reported within this paper.

## Author Contributions

Conceptualization: KLB, ARL

Data curation: KLB, TS, LP

Formal Analysis: KLB, TS

Funding acquisition: ARL

Methodology: KLB, TS, ARL

Project administration: KLB, ARL

Resources: ARL, EGJ

Software: KLB, TS

Supervision: MMH, ARL

Visualization: KLB

Writing – original draft: KLB, ARL

Writing – review & editing: KLB, TS, LP, CT, EGJ, MMH, ARL

