## Supplementary Material for "Idiosyncrasy and generalizability of contraceptive- and hormone-related functional connectomes across the menstrual cycle"

This document includes:

- Supplemental Methods
- Supplemental Table 1
- Supplemental Figures 1-3

### Supplemental Methods

#### Anatomical Data Preprocessing

A total of 1 T1-weighted (T1w) images were found within the input BIDS dataset. The T1-weighted (T1w) image was corrected for intensity non-uniformity (INU) with N4BiasFieldCorrection (Tustison et al. 2010), distributed with ANTs 2.3.3 (Avants et al. 2008, RRID:SCR_004757), and used as T1w-reference throughout the workflow. The T1w-reference was then skull-stripped with a Nipype implementation of the antsBrainExtraction.sh workflow (from ANTs), using OASIS30ANTs as target template. Brain tissue segmentation of cerebrospinal fluid (CSF), white-matter (WM) and gray-matter (GM) was performed on the brain-extracted T1w using fast (FSL 5.0.9, RRID:SCR_002823, Zhang, Brady, and Smith 2001). Brain surfaces were reconstructed using recon-all (FreeSurfer 6.0.1, RRID:SCR_001847, Dale, Fischl, and Sereno 1999), and the brain mask estimated previously was refined with a custom variation of the method to reconcile ANTs-derived and FreeSurfer-derived segmentations of the cortical gray-matter of Mindboggle (RRID:SCR_002438, Klein et al. 2017). Volume-based spatial normalization to one standard space (MNI152NLin2009cAsym) was performed through nonlinear registration with antsRegistration (ANTs 2.3.3), using brain-extracted versions of both T1w reference and the T1w template. The following template was selected for spatial normalization: ICBM 152 Nonlinear Asymmetrical template version 2009c [Fonov et al. (2009), RRID:SCR_008796; TemplateFlow ID: MNI152NLin2009cAsym].

#### Single-Echo Functional Data Preprocessing

For each of the 60 BOLD runs found per subject (across all tasks and sessions), the following preprocessing was performed. First, a reference volume and its skull-stripped version were generated using a custom methodology of *fMRIPrep*. Susceptibility distortion correction (SDC) was omitted. The BOLD reference was then co-registered to the T1w reference using `bbregister` (FreeSurfer) which implements boundary-based registration (Greve and Fischl 2009). Co-registration was configured with six degrees of freedom. Head-motion parameters with respect to the BOLD reference (transformation matrices, and six corresponding rotation and translation parameters) are estimated before any spatiotemporal filtering using `mcflirt` (FSL 5.0.9, Jenkinson et al. 2002). BOLD runs were slice-time corrected using `3dTshift` from AFNI 20160207 (Cox and Hyde 1997, RRID:SCR_005927). The BOLD time-series were resampled onto the following surfaces (FreeSurfer reconstruction nomenclature): *fsnative*, *fsaverage5*. The BOLD time-series (including slice-timing correction when applied) were resampled onto their original, native space by applying the transforms to correct for head-motion. These resampled BOLD time-series will be referred to as *preprocessed BOLD in original space*, or just *preprocessed BOLD*. The BOLD time-series were resampled into standard space, generating a *preprocessed BOLD run in MNI152NLin2009cAsym space*. First, a reference volume and its skull-stripped version were generated using a custom methodology of *fMRIPrep*. Several confounding time-series were calculated based on the *preprocessed BOLD*: framewise displacement (FD), DVARS and three region-wise global signals. FD was computed using two formulations following Power (absolute sum of relative motions, Power et al. (2014)) and Jenkinson (relative root mean square displacement between affines, Jenkinson et al. (2002)). FD and DVARS are calculated for each functional run, both using their implementations in *Nipype* (following the definitions by Power et al. 2014). The three global signals are extracted within the CSF, the WM, and the whole-brain masks. Additionally, a set of physiological regressors were extracted to allow for component-based noise correction (*CompCor*, Behzadi et al. 2007). Principal components are estimated after high-pass filtering the *preprocessed BOLD* time-series (using a discrete cosine filter with 128s cut-off) for the two *CompCor* variants: temporal (tCompCor) and anatomical (aCompCor). tCompCor components are then calculated from the top 2% variable voxels within the brain mask. For aCompCor, three probabilistic masks (CSF, WM and combined CSF+WM) are generated in anatomical space. The implementation differs from that of Behzadi et al. in that instead of eroding the masks by 2 pixels on BOLD space, the aCompCor masks are subtracted a mask of pixels that likely contain a volume fraction of GM. This mask is obtained by dilating a GM mask extracted from the FreeSurfer's *aseg* segmentation, and it ensures components are not extracted from voxels containing a minimal fraction of GM. Finally, these masks are resampled into BOLD space and binarized by thresholding at 0.99 (as in the original implementation). Components are also calculated separately within the WM and CSF masks. For each CompCor decomposition, the *k* components with the largest singular values are retained, such that the retained components' time series are sufficient to explain 50 percent of variance across the nuisance mask (CSF, WM, combined, or temporal). The remaining components are dropped from consideration. The head-motion estimates calculated in the correction step were also placed within the corresponding confounds file. The confound time series derived from head motion estimates and global signals were expanded with the inclusion of temporal derivatives and quadratic terms for each (Satterthwaite et al. 2013). Frames that exceeded a threshold of 0.5 mm FD or 1.5 standardised DVARS were annotated as motion outliers. All resamplings can be performed with *a single interpolation step* by composing all the pertinent transformations (i.e. head-motion transform matrices, susceptibility distortion correction when available, and co-registrations to anatomical and output spaces). Gridded (volumetric) resamplings were performed using `antsApplyTransforms` (ANTs), configured with Lanczos interpolation to minimize the smoothing effects of other kernels (Lanczos 1964). Non-gridded (surface) resamplings were performed using `mri_vol2surf` (FreeSurfer).

Many internal operations of *fMRIPrep* use *Nilearn* 0.6.2 (Abraham et al. 2014, RRID:SCR_001362), mostly within the functional processing workflow. For more details of the pipeline, see the section corresponding to workflows in *fMRIPrep*'s documentation.

**Multi-Echo Functional Data Preprocessing**

For each of the BOLD runs per subject, the following preprocessing was performed. First, a reference volume and its skull-stripped version were generated by aligning and averaging the first echo of 4 single-band references (SBRefs). A B0-nonuniformity map (or fieldmap) was estimated based on two (or more) echo-planar imaging (EPI) references with opposing phase-encoding directions, with 3dQwarp Cox and Hyde (1997) (AFNI 20160207). Based on the estimated susceptibility distortion, a corrected EPI (echo-planar imaging) reference was calculated for a more accurate co-registration with the anatomical reference. The BOLD reference was then co-registered to the T1w reference using bbregister (FreeSurfer) which implements boundary-based registration (Greve and Fischl 2009). Co-registration was configured with six degrees of freedom. Head-motion parameters with respect to the BOLD reference (transformation matrices, and six corresponding rotation and translation parameters) are estimated before any spatiotemporal filtering using mcflirt (FSL 5.0.9, Jenkinson et al. 2002). BOLD runs were slice-time corrected using 3dTshift from AFNI 20160207 (Cox and Hyde 1997, RRID:SCR_005927). The BOLD time-series (including slice-timing correction when applied) were resampled onto their original, native space by applying a single, composite transform to correct for head-motion and susceptibility distortions. These resampled BOLD time-series will be referred to as preprocessed BOLD in original space, or just preprocessed BOLD. A T2* map was estimated from the preprocessed BOLD by fitting to a monoexponential signal decay model with nonlinear regression, using T2*/S0 estimates from a log-linear regression fit as initial values. For each voxel, the maximal number of echoes with reliable signal in that voxel were used to fit the model. The calculated T2* map was then used to optimally combine preprocessed BOLD across echoes following the method described in (Posse et al. 1999). The optimally combined time series was carried forward as the preprocessed BOLD. First, a reference volume and its skull-stripped version were generated using a custom methodology of fMRIPrep. The BOLD time-series were resampled onto the following surfaces (FreeSurfer reconstruction nomenclature): fsnative, fsaverage5. The BOLD time-series were resampled into standard space, generating a preprocessed BOLD run in MNI152NLin2009cAsym space. First, a reference volume and its skull-stripped version were generated using a custom methodology of fMRIPrep. Several confounding time-series were calculated based on the preprocessed BOLD: framewise displacement (FD), DVARS and three region-wise global signals. FD was computed using two formulations following Power (absolute sum of relative motions, Power et al. (2014)) and Jenkinson (relative root mean square displacement between affines, Jenkinson et al. (2002)). FD and DVARS are calculated for each functional run, both using their implementations in Nipype (following the definitions by Power et al. 2014). The three global signals are extracted within the CSF, the WM, and the whole-brain masks. Additionally, a set of physiological regressors were extracted to allow for component-based noise correction (CompCor, Behzadi et al. 2007). Principal components are estimated after high-pass filtering the preprocessed BOLD time-series (using a discrete cosine filter with 128s cut-off) for the two CompCor variants: temporal (tCompCor) and anatomical (aCompCor). tCompCor components are then calculated from the top 2% variable voxels within the brain mask. For aCompCor, three probabilistic masks (CSF, WM and combined CSF+WM) are generated in anatomical space. The implementation differs from that of Behzadi et al. in that instead of eroding the masks by 2 pixels on BOLD space, the aCompCor masks are subtracted from a mask of pixels that likely contain a volume fraction of GM. This mask is obtained by dilating a GM mask extracted from the FreeSurfer's aseg segmentation, and it ensures components are not extracted from voxels containing a minimal fraction of GM. Finally, these masks are resampled into BOLD space and binarized by thresholding at 0.99 (as in the original implementation). Components are also calculated separately within the WM and CSF masks. For each CompCor decomposition, the k components with the largest singular values are retained, such that the retained components' time series are sufficient to explain 50 percent of variance across the nuisance mask (CSF, WM, combined, or temporal). The remaining components are dropped from consideration. The head-motion estimates calculated in the correction step were also placed within the corresponding confounds file. The confound time series derived from head motion estimates and global signals were expanded with the inclusion of temporal derivatives and quadratic terms for each (Satterthwaite et al. 2013). Frames that exceeded a threshold of 0.5 mm FD or 1.5 standardised DVARS were annotated as motion outliers. All resamplings can be performed with a single interpolation step by composing all the pertinent transformations (i.e. head-motion transform matrices, susceptibility distortion correction when available, and co-registrations to anatomical and output spaces). Gridded (volumetric) resamplings were performed using antsApplyTransforms (ANTs), configured with Lanczos interpolation to minimize the smoothing effects of other kernels (Lanczos 1964). Non-gridded (surface) resamplings were performed using mri_vol2surf (FreeSurfer).

Many internal operations of fMRIPrep use Nilearn 0.6.2 (Abraham et al. 2014, RRID:SCR_001362), mostly within the functional processing workflow. For more details of the pipeline, see the section corresponding to workflows in fMRIPrep's documentation.

**Copyright Waiver**

The above boilerplate text was automatically generated by fMRIPrep with the express intention that users should copy and paste this text into their manuscripts unchanged. It is released under the CC0 license.

### Supplemental Results

Pairwise product-moment and point biserial (for dichotomous variables site and HC use) correlation coefficients and associated significance testing revealed significant associations between non-brain variables in this dataset. Dataset was significantly associated with hormonal contraceptive (HC) use and menstrual cycle day was significantly associated with [$E_2$] ($p < 0.01$). Abbreviations: $E_2$ = estradiol; $P_4$ = progesterone

**Supplementary Table 1. Correlations between hormone, lifestyle, and logistical variables**

| | Dataset | HC Use | [$E_2$] | [$P_4$] | Head Motion |
|---|---|---|---|---|---|
| Dataset | -- | 4x10$^{-9}$ | 1 | 1 | 0.00 |
| HC Use | ***0.71*** | -- | 0.093 | 0.046 | 0.82 |
| [$E_2$] | 0 | -0.24 | -- | 0.15 | 0.32 |
| [$P_4$] | 0 | *-0.29* | 0.21 | -- | 0.65 |
| Head motion | ***0.67*** | 0.027 | -0.12 | 0.06 | – |

*Note.* Values above the diagonal are *p*-values to the corresponding Pearson's correlation coefficients shown below the diagonal. Significant correlations at p < 0.05 are indicated in italics, while significant correlations at p < 0.01 are indicated in bold italics.

***Supplementary Table 2*. Performance and generalizability of contraceptive- and hormone-related subnetworks: Sensitivity analysis without high-$E_2$ days.**

| | HC use | Estradiol | | Progesterone | |
|---|---|---|---|---|---|
| | | Base | HC use | Base | HC use |
| CV mean ± sd | 79.9% ± 11.1% | 0.097 ± 0.312 | 0.043 ± 0.336 | 0.285 ± 0.277 | 0.236 ± 0.283 |
| Alpha | 0.0001 | 0.359 | 2.783 | 21.544 | $1.00 \times 10^5$ |
| In-sample (28and*) | | | | | |
| Model performance | 80% | 0.753 | 0.968 | 0.946 | 0.653 |
| MSE | 0.1833 | 0.350 | 0.024 | 0.143 | 0.515 |
| Out-of-sample (DIVA) | | | | | |
| Model performance | 50% | 0.103 | 0.054 | 0.571 | 0.357 |
| MSE | 0.5 | 0.960 | 0.591 | 0.148 | 0.094 |

*Note*. HC use models are logistic regression with $L_2$ penalties and their performance is percent accuracy of HC use predicted by functional connectivity. Estradiol and progesterone models are linear regression with $L_2$ penalties (i.e., Ridge regression) and their performance is provided by the Spearman correlation between true hormone concentrations and hormone concentrations predicted from functional connectivity of the identified subnetwork.

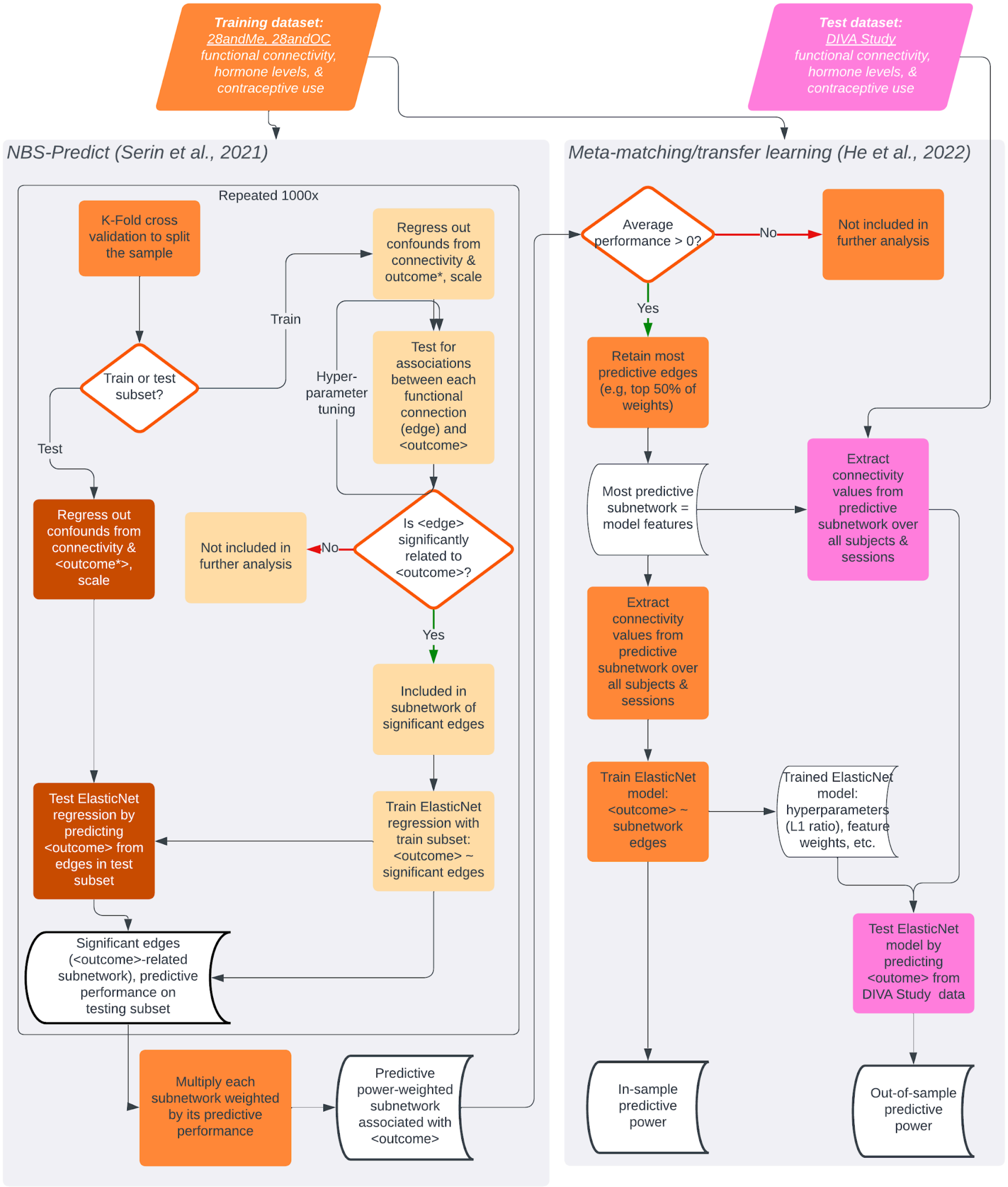


***Supplementary Figure 1.*** **NBS-Predict and transfer learning to identify generalizable functional subnetworks associated with HC use and hormone levels.**
Pipeline was run to build models of functional subnetworks related to each HC use, estradiol [$E_2$], and progesterone [$P_4$], separately. Orange shapes indicate processes with 28andMe and 28andOC data, pink shapes indicate processes with DIVA Study data. Logistic regression with $L_2$ penalties (i.e., Ridge) was used for HC use, and performance was assessed as classification accuracy. Linear regressions with $L_2$ penalties were used for [$E_2$] and [$P_4$] and performance was assessed with Spearman correlations.

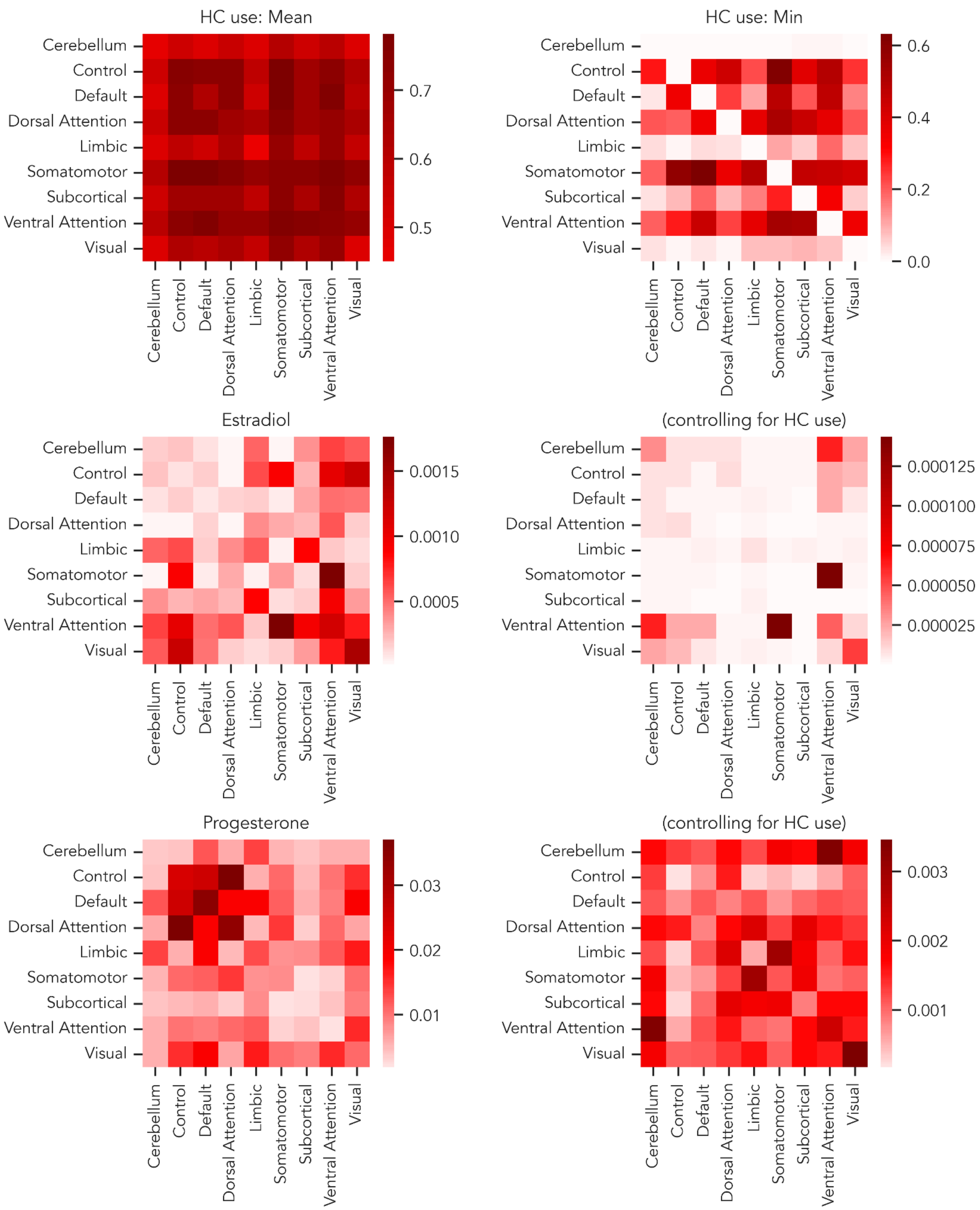


***Supplementary Figure 2.*** **Network-wise mean relatedness of regional connectivity to HC use (top row), estradiol fluctuations (middle row), and progesterone fluctuations (bottom row).** In the case of estradiol and progesterone, the left column shows mean relatedness without accounting for HC use and the right column shows mean relatedness after regressing out HC use. Top right graph shows the average *minimum* relatedness per large-scale brain network, due to relative homogeneity in the average relatedness.

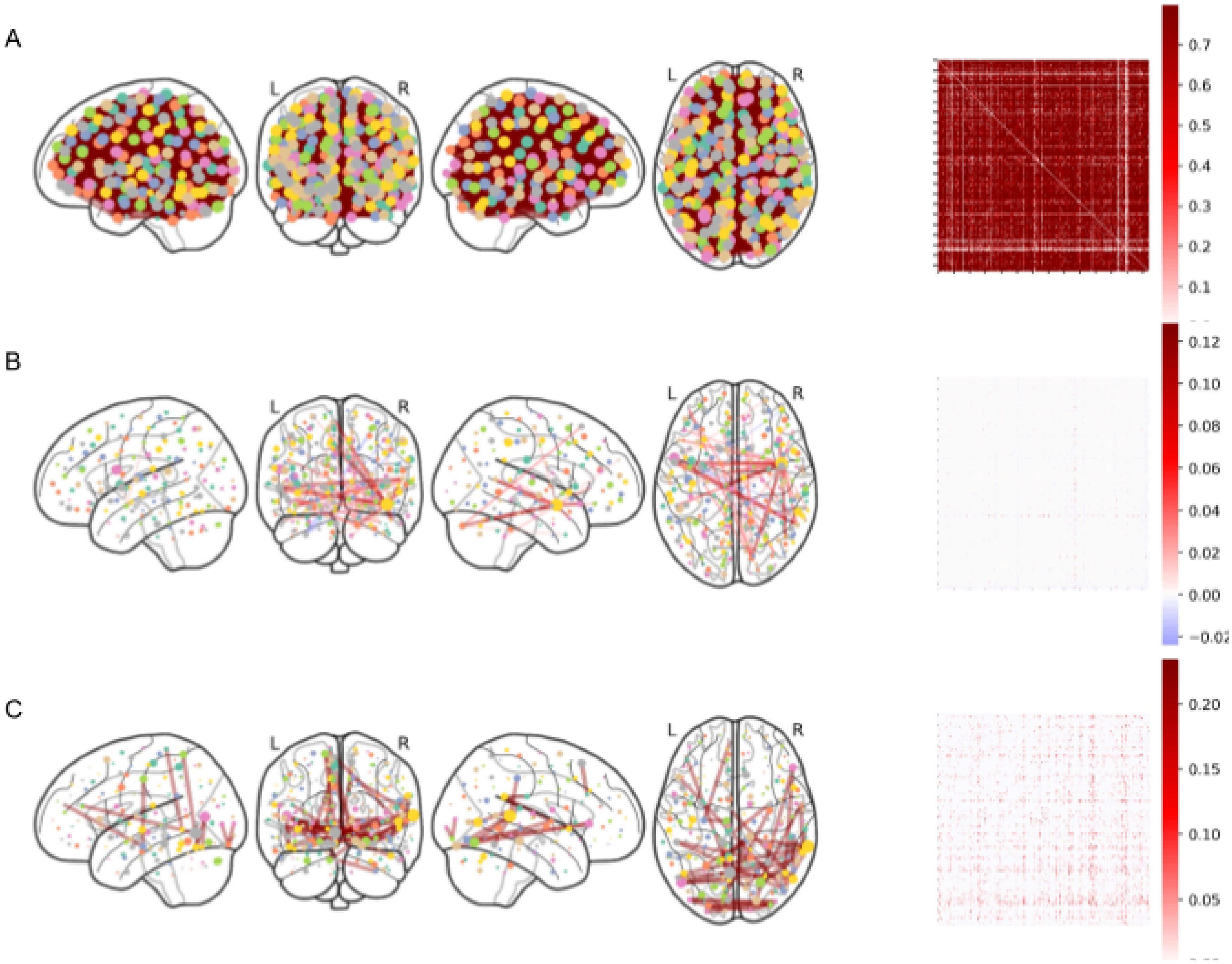


***Supplementary Figure 3. Contraceptive- and hormone-related functional subnetworks, identified without data from the ovulatory window.*** (A) HC-related subnetwork is virtually identical regardless of the inclusion of high-$E_2$ data from the ovulatory window, as are (B) [$E_2$]- and (C) [$P_4$]-related subnetworks.Edges in the glass brain plots and accompanying correlograms reflect the average contraceptive- and hormone-related connections, weighted by model performance (i.e., accuracy in (A), Spearman correlations in (B, C)).
